# Effects of a postnatal *Atrx* conditional knockout in neurons on autism-like behaviours in male and female mice

**DOI:** 10.1101/2020.02.05.936260

**Authors:** Nicole Martin-Kenny, Nathalie G. Bérubé

**Author notes:** Corresponding author at Western University, London, Ontario, Canada.

## Abstract

**Background:** Alpha-thalassemia/mental retardation, X-linked, or *ATRX*, is an autism susceptibility gene that encodes a chromatin remodeler. Mutations of *ATRX* result in the ATR-X intellectual disability syndrome and have been identified in autism spectrum disorder (ASD) patients. The mechanisms by which *ATRX* mutations lead to autism and autistic-like behaviours are not yet known. To address this question, we generated mice with postnatal *Atrx* inactivation in excitatory neurons of the forebrain and performed a battery of behavioural assays that assess autistic-like behaviours.

**Methods:** Male and female mice with a postnatal conditional knockout of *Atrx* were tested in a battery of behavioural tests that assess autistic features. We utilized paradigms that measure social behaviour, repetitive and stereotyped behaviours, as well as sensory gating. Statistics were calculated by two-way repeated measures ANOVA with Sidak’s multiple comparison test or unpaired Student’s T-tests as indicated.

**Results:** The behaviour tests revealed no significant differences between *Atrx*-cKO and control mice. We identified sexually dimorphic changes in odor habituation and discrimination; however, these changes did not correlate with social deficits. We additionally observed sex-specific differences in sociability, vertical episodes, and acoustic startle response when results were analyzed by sex.

**Conclusion:** The postnatal knockout of *Atrx* in forebrain excitatory neurons does not lead to autism-related behaviours in male or female mice.

## Background

Autism spectrum disorder (ASD) is a behaviourally defined condition characterized by deficits in social and communicative abilities, impaired sensory gating, as well as the presence of stereotyped behaviours (1,2). Recent work has highlighted the important contribution of *de novo* variants and inherited copy number variants in ASD, confirming a strong genetic component of this disease (1–3). Numerous autism susceptibility genes have been identified and shown to share commonalities in synaptic, transcriptional, and epigenetic mechanisms (4–6). Mouse models have typically been used to investigate the behavioural implications of genetic mutations associated with ASD (7,8). However, these studies often omit the investigation of the sex-specific effects of these genetic mutations, limiting the potential translational applications. In the general population, ASD occurs at a 4:1 male:female ratio, highlighting the need to study the outcome of genetic mutations in both male and female model systems (1,2).

In this study, we describe the impact of targeted inactivation of *Atrx* in glutamatergic neurons on behaviours related to autism in male and female mice. ATRX belongs to the SWI/SNF family of chromatin remodeling factors (9,10). Mutations in the *ATRX* gene are associated with an intellectual disability syndrome referred to as ATRX-syndrome, characterized by autistic-like behaviours in addition to cognitive deficits, intellectual disabilities, and developmental delays (11). Furthermore, autistic carriers of a rare mutation in *ATRX* have been discovered and missense variants in *ATRX* have been identified in male ASD patients. Interestingly, female carriers of *ATRX* mutations experience skewed X-inactivation, and as a result, are asymptomatic (12–18).

Previous studies have demonstrated that the loss of *Atrx* in mouse forebrain causes changes in gene expression (19,20). Specifically, transcription of autism susceptibility genes, including the monogenic *Neuroligin-4* (*Nlgn4)*, are altered upon loss of *Atrx* (19). Additionally, sexually dimorphic transcript changes have been revealed in the adult mouse hippocampus upon the loss of *Atrx* in excitatory neurons(21). However, autistic behaviours were not evaluated in that report, and should be addressed given the link between *ATRX* mutations and ASD.

In this study, we characterize the impact of *Atrx* loss in neurons on autistic-like behaviours in male and female mice. We used the AtrxCamKIICre model (21) where *Atrx* is ablated in forebrain excitatory neurons postnatally, thus bypassing deleterious effects of ATRX loss-of-function previously observed in neural progenitors during brain development (22,23). An array of behavioural assays was performed to investigate the presence of autistic-like behaviours, including deficits in sociability, altered sensory gating, and the presence of repetitive or stereotyped behaviours. These investigations revealed minimal behavioural deficits related to autism in both male and female mice. We did, however, identify sex-specific differences in sociability, vertical episode stereotypies, and startle responses when behavioural analyses were grouped by sex. Overall, this study demonstrates that a conditional loss of *Atrx* in forebrain excitatory neurons postnatally does not result in autistic-like behaviours in male or female mice.

## Materials and Methods

### Animal Care and husbandry

Mice were exposed to a 12-hour-light/12-hour-dark cycle and with water and chow ad libitum. The *Atrx*^loxP^ mice have been described previously(23). *Atrx*^loxP^ mice were mated with C57BL/6 mice expressing Cre recombinase under the control of the *αCaMKII* gene promoter(24). The progeny includes hemizygous male mice that produce no ATRX protein in forebrain excitatory neurons (*Atrx*-cKO^MALE^). The *Atrx*-cKO males were mated to *Atrx*^loxP^ females to yield homozygous deletion of *Atrx* in female mice (*Atrx*-cKO^FEMALE^). Male and female littermate floxed mice lacking the Cre allele were used as controls (Ctrl^MALE^; Ctrl^FEMALE^). Genotyping of tail biopsies for the presence of the floxed and Cre alleles was performed as described previously(23). All procedures involving animals were conducted in accordance with the regulations of the Animals for Research Act of the province of Ontario and approved by the University of Western Ontario Animal Care and Use Committee (2017-048). Behavioural assessments started with less demanding tasks and moved to more demanding tasks in the following order: open field test, marble-burying assay, induced self-grooming, pre-pulse inhibition (PPI) and startle response, social approach, and 3-chamber social tests. ARRIVE guidelines were followed: mouse groups were randomized, experimenters were blind to the genotypes, and software-based analysis was used to score mouse performance in all the tasks. All behavioural tasks were performed between 9:00 AM and 4:00 PM. All behavioural assays were performed when mice were between 3-7 months of age. 3 cohorts of male and female mice were used to reach the final sample size (Ctrl^MALE^: 17; *Atrx*-cKO^MALE^: 10; Ctrl^FEMALE^:13; *Atrx*-cKO^FEMALE^: 13). Statistics were calculated by two-way repeated measures ANOVA with Sidak’s multiple comparison test or unpaired Student’s T-tests as indicated in the figure legends.

### Odor Habituation and Discrimination

The odor habituation and discrimination assay was performed as previously described(25) to assess olfaction. Individual mice were placed into a clean cage with a wire lid and allowed to habituate to the testing room for 30 minutes. The mice were then presented with an odor on a cotton swab (either almond, banana, or water as a control) for a 2-minute trial. For each trial, 50µl of water, almond extract, or banana extract (Club House) was pipetted onto the tip of a cotton swab and the swab was then secured to the wire cage top through the water bottle opening. The mice were presented with the same odor three times before being presented with a new odour, for a total of nine trials. During the 2-minute trials, the amount of time that the mouse spent sniffing the odor was recorded by an investigator blind to the genotype. Sniffing was defined as the animal’s nose being in proximity to the cotton swab (2cm or closer), and oriented toward the swab.

### Social Approach

This test was performed as previously described(26) to assess for sociability with conspecific mice. For two consecutive days prior to the test day, individual mice were habituated to the open area for 10 minutes. On the test day, pairs of unfamiliar, same-sex conspecific mice were placed into the cage. Behaviour of the mice was recorded by the AnyMaze software and video-tracking system. The time spent in social interaction, defined as the experimental mouse sniffing the stranger mouse, was manually scored by investigators unaware of the genotype.

### 3-Chamber Social Tests (Social Preference and Novelty)

The social preference and social novelty assessments were performed as described(26,27) with minor modifications. Individual mice were placed in the 3-chambered box and allowed to freely explore the arena during a 10-minute habituation period. After the habituation period, an unfamiliar, same-sex mouse of a different genotype (stranger 1) was placed in one of the side chambers under a wire cage. An identical wire cage containing an inanimate object was placed in the opposite chamber. The test mouse was then allowed to explore the entire 3-chambered arena for 10 minutes. The amount of time spent in each chamber was recorded by the AnyMaze video-tracking system. Following this period, a second unfamiliar, same-sex mouse of a different genotype (stranger 2) was placed into the wire cage previously containing the inanimate object. The test mouse was then allowed to explore the 3-chambered arena for 10 minutes. The amount of time spent in each chamber was recorded by the AnyMaze video-tracking system. Based on the amount of time spent in each chamber, a ‘sociability index’ and a ‘social novelty index’ was calculated as previously described(27). The sociability index was calculated as: time_stranger_/(time_stranger_+time_object_) × 100. The social novelty index was calculated as: time_novel_/(time_novel_+time_familiar_) × 100.

### Marble Burying

The test was performed as previously described(28) with modifications to evaluate repetitive digging behaviour. Mice were brought into the test room to habituate in their home cages for approximately 30 minutes prior to the test. The test cages were filled with 4cm of wood-chip bedding, with 12 evenly spaced glass marbles placed on the surface. Individual mice were then placed in the test cage and permitted to explore for 30 minutes. Following the test, the number of marbles buried (>3/4 surface covered) was counted and recorded by investigators blind to the genotype.

### Induced Self-Grooming

The test was performed as previously described (27,29) to evaluate repetitive grooming tendencies. Mice were individually habituated in an empty test cage for 30 minutes prior to the test. To amplify natural grooming tendencies, mice were misted with water 3 times at 10cm distance of the upper-back. Following this misting, the grooming behaviour of each mouse was recorded by the Anymaze video-tracking system for 30 minutes. The time that each individual mouse spent grooming during this 30-minute trial was manually scored by the rater, unaware of the genotype.

### Open Field Test

Mice were brought into the testing room to habituate in their home cages approximately 30 minutes prior to the test. Mice were placed in a 20 cm × 20 cm arena with 30 cm high walls. Locomotor activity was automatically recorded in 5-minute intervals over 2 hours (AccuScanInstrument)(30). For each mouse the number of vertical episodes was assessed.

### Pre-Pulse Inhibition of the Startle Response

The pre-pulse inhibition and startle response tests were performed as previously described(31) to assess sensory gating. Mice underwent two days of habituation prior to the testing day, to acclimate the mice to the apparatus. During this habituation, mice were individually placed in the chamber apparatus and exposed to background noise (65 db) for 5 minutes (SR-LAB, San Diego Instruments). On the test day, individual mice were placed in the chamber and acclimated for 10 minutes with background noise. The mice then underwent a habituation block, consisting of fifty acoustic startle trials, with 20 ms stimulus of 115 db, and intertrial interval of 20 seconds. After the habituation block, mice underwent a prepulse-inhibition block consisting of ten sets of five types of trials randomly ordered with variable intertrial intervals of 10, 15, or 20 seconds. Four of the five trial types consisted of prepulses (intensity of 75 or 80 db, length of 20 ms), separated from the startle stimulus (intensity of 115 db, length of 40 ms) by an interstimulus interval of either 30 ms or 100 ms. The fifth trial type was a startle pulse alone. The startle response was measured by the movement of the mouse on the platform, which generates a transient force analyzed by the software. The startle magnitude recorded was an average for the ten trials of each trial type and startle magnitudes of pre-pulse trials were normalized to the pulse-only trial.

## Results

### Sexually dimorphic olfaction defects in Atrx-cKO mice

As olfactory impairments can confound the interpretation of other tests, especially social behaviour assays, we first wanted to address whether the loss of *Atrx* in excitatory neurons of the forebrain alters olfaction in male and female mice. To do this, we performed the odor discrimination and habituation assay (25). In this test, mice were presented with multiple odors for 2-minute trials, during which the amount of time spent sniffing the odor was recorded. During this test, *Atrx-*cKO^MALE^ mice spent significantly less time sniffing the odors throughout the nine trials compared to Ctrl^MALE^ mice (ANOVA, **p=0.004; **Fig.1A**). In particular, *Atrx-*cKO^MALE^ mice spent significantly less time sniffing the cotton swab when first presented with the banana odor (multiple comparisons, ***p<0.001). There was no significant difference in the overall amount of time spent sniffing the odors throughout the test between *Atrx-*cKO^FEMALE^ and Ctrl^FEMALE^ mice. However, *Atrx-*cKO^FEMALE^ mice did spend significantly more time sniffing the cotton swab when first presented with the banana odor (multiple comparison, ****p<0.0001; Fig. 1B). Overall, the results of this test suggest that the loss of *Atrx* in forebrain excitatory neurons results in sexually dimorphic changes in olfaction that must be considered in subsequent behaviour testing of these mice.

**Figure 1.**
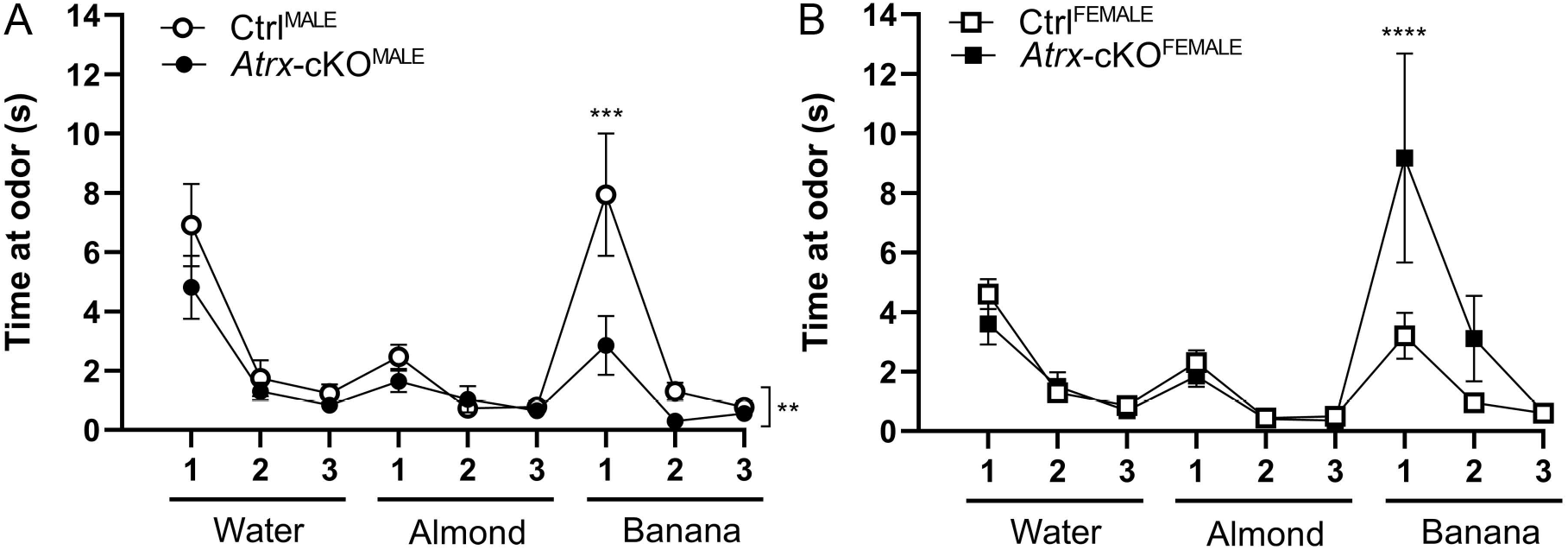
*Atrx*-cKO^MALE^ and *Atrx*-cKO^FEMALE^ mice exhibit differences in olfaction during the odor habituation and discrimination assay. The amount of time spent sniffing a cotton swab saturated with an odor during nine, 2-minute trials. A) *Atrx*-cKO^MALE^ (n=24) mice spend less time sniffing cotton swabs with corresponding odors compared to Ctrl^MALE^ (n=17) (twANOVA, F_(1,351)_=8.203, **p=0.004; mc, ***p<0.001). B) *Atrx*-cKO^FEMALE^ (n=19) mice spend more time sniffing the cotton swab with the banana odor when first exposed to the scent compared to Ctrl^FEMALE^ (n=15) (twANOVA, F_(1,288)_=3.188, p=0.075; mc, ****p<0.0001). Error bars: ± SEM.

### Social assays reveal sex-differences, but not genotypic-differences, in Atrx-cKO^MALE^ and Atrx-cKO^FEMALE^ mice

Given that *ATRX* mutations are associated with autistic traits in humans, we next sought to investigate if the loss of *Atrx* in forebrain excitatory neurons has any effect on social behaviour. Changes in sociability and social preference are some of the most common deficits observed in mouse models with autism-associated genetic mutations (26,27,32–34). As such, we first investigated sociability of *Atrx-*cKO^MALE^ and *Atrx-*cKO^FEMALE^ mice by means of the social approach assay, as described previously (26). There was no significant difference in the total amount of time that *Atrx-*cKO^MALE^ and *Atrx-*cKO^FEMALE^ mice spent interacting with a stranger mouse compared to controls (Fig. 2A). However, when these results were grouped and analyzed by sex, male mice (*Atrx-*cKO^MALE^ and Ctrl^MALE^) spent more time socially interacting with the stranger mouse compared to female mice (*Atrx-*cKO^FEMALE^ and Ctrl^FEMALE^) (ANOVA, *p=0.048; Fig 2A). When social interaction was analyzed over one-minute intervals during the 10-minute test, there were no genotypic or sex-differences (Fig. 2B).

**Figure 2.**
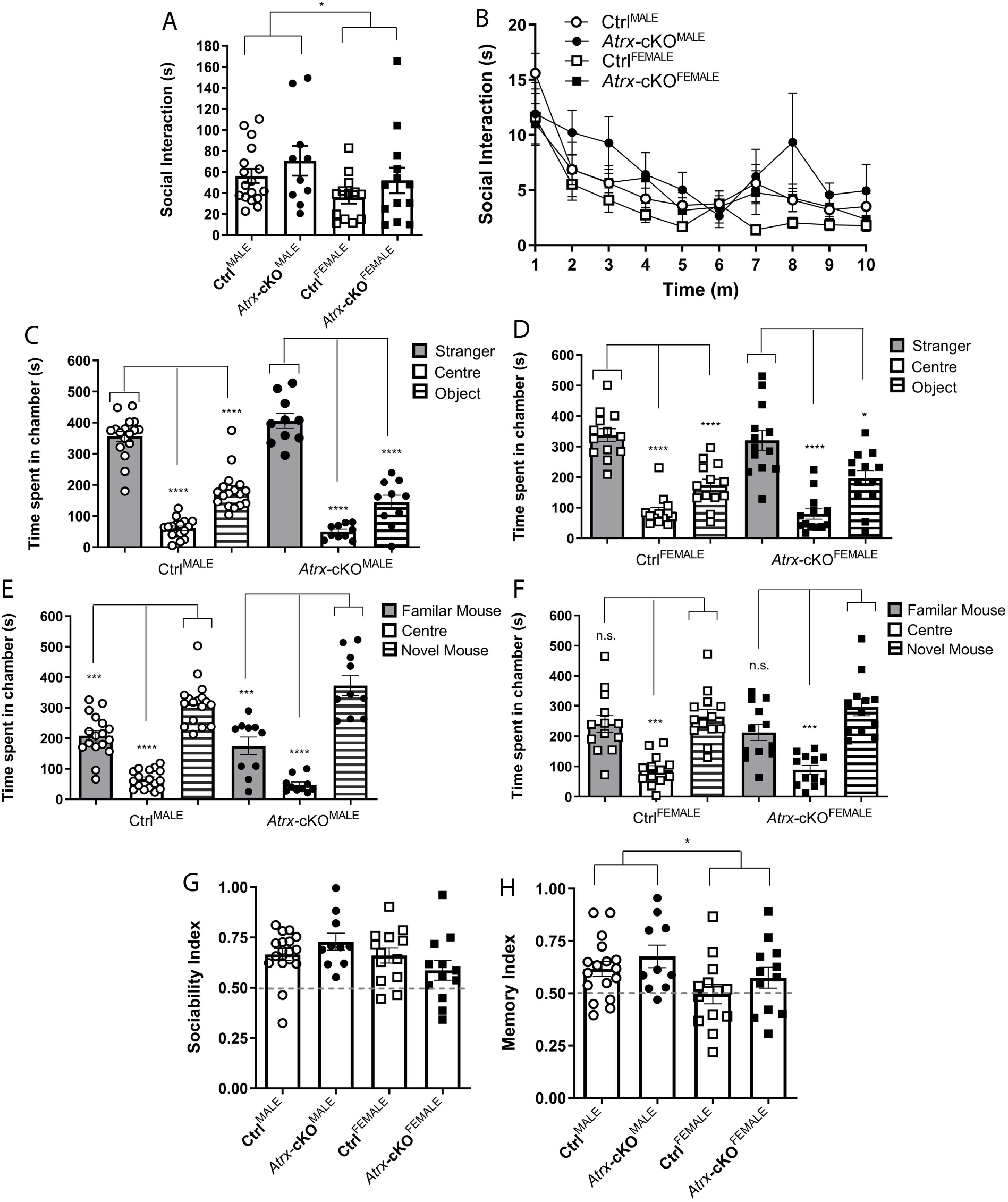
Social behaviour assays reveal sex-differences in sociability, despite lack of genotypic difference between *Atrx*-cKO and control mice. A) Total amount of social interaction with a mouse conspecific for genotype and sex during the 10-minute social approach assay (twANOVA, genotype, F_(1,49)_=2.496, p=0.121; twANOVA, sex, F_(1,49)_=4.086, *p=0.048). B) Social interaction over 1-minute intervals during the social approach assay (twANOVA, genotype, F_(1,51)_=1.489, p=0.228; twANOVA, sex, F_(1,51)_=3.425, p=0.070). C) Amount of time Ctrl^MALE^(twANOVA, F_(2,32)_=78.39, ****p<0.0001; mc, ****p<0.0001) and *Atrx*-cKO^MALE^ (twANOVA, F_(2,18)_=58.26, ****p<0.0001; mc, ****p<0.0001) spent in the empty centre chamber and chambers containing either a stranger mouse or novel object in the social preference assay. D) Amount of time Ctrl^FEMALE^ (twANOVA, F_(2,24)_=31.39, ****p<0.0001; mc, ****p<0.0001) and *Atrx*-cKO^FEMALE^ (twANOVA, F_(2,24)_=14.88, ****p<0.0001; mc, *p<0.05, ****p<0.0001) spent in chambers containing either a stranger mouse or novel object, or the empty centre chamber E) Time spent in the empty centre chamber and chambers containing a familiar mouse or a novel mouse in the social novelty assay for E) Ctrl^MALE^ (twANOVA, F_(2,32)_=51.38, ****p<0.0001; mc, ***p<0.001, ****p<0.0001) and *Atrx*-cKO^MALE^ mice (twANOVA, F_(2,18)_=27.26, ****p<0.0001; mc, ***p<0.001, ****p<0.0001), and F) Ctrl^FEMALE^ (twANOVA, F_(2,24)_=11.77, ***p<0.001; mc, ***p<0.001) and *Atrx*-cKO^FEMALE^ mice (twANOVA, F_(2,22)_=12.50, ***p<0.001; mc, ***p<0.001). G) The sociability index for each mouse was calculated as the time spent in the stranger mouse chamber, divided by the total time in the stranger mouse and novel object chambers (twANOVA, genotype, F_(1,28)_=0.019, p=0.890; twANOVA, sex, F_(1,48)_=3.574, p=0.065). H) The social memory index for each mouse was calculated as the time spent in the novel mouse chamber, divided by the total time in the familiar mouse and novel mouse chambers (twANOVA, genotype, F_(1,48)_=2.217, p=0.143; twANOVA, sex, F_(1,48)_=5.811, *p=0.019). *Atrx*-cKO^MALE^: n=10, Ctrl^MALE^: n=17, *Atrx*-cKO^FEMALE^: n=13, Ctrl^FEMALE^: n=13. Error bars: ± SEM.

We also investigated social preference and social novelty in the three-chambered paradigm (27). During the first part of the paradigm, social preference was assessed as mice were placed into a three-chambered apparatus and were free to explore between the chambers. The outer two chambers contained either a novel object or a stranger mouse, while the centre chamber remained empty. Both Ctrl^MALE^ and *Atrx-*cKO^MALE^ mice demonstrated a preference for the chamber containing a stranger mouse compared to the object and the empty chamber (multiple comparisons, ****p<0.0001; Fig. 2C). Similarly, both Ctrl^FEMALE^ and *Atrx-*cKO^FEMALE^ mice preferred exploration of the stranger mouse (multiple comparisons, *p=0.030, ****p<0.0001; Fig. 2D). No genotypic differences were observed in social preference between groups (Fig. 2C-D).

Social novelty was investigated during the second part of the paradigm in which the outer chambers contained either the stranger mouse from the first part of the test (familiar mouse) or a novel mouse. Ctrl^MALE^ and *Atrx-*cKO^MALE^ mice both spent more time in the chamber containing the novel mouse compared to the familiar mouse and the empty chamber (multiple comparisons, ***p<0.001, ****p<0.0001; Fig. 2E). Interestingly, although Ctrl^FEMALE^ and *Atrx-*cKO^FEMALE^ mice both spent significantly less time in the empty chamber, neither demonstrate a preference for the novel mouse over the familiar mouse (multiple comparisons, ***p<0.001; Fig. 2F). Sociability and social memory indexes were calculated based on the social preference and social novelty results. *Atrx-*cKO and Ctrl mice did not display genotypic or sex-differences in their sociability indexes (Fig. 2G). Similarly, there was no genotypic difference in social memory indexes, however, male mice (Ctrl^MALE^ and *Atrx-*cKO^MALE^) display a greater social memory index than female mice (Ctrl^FEMALE^ and *Atrx-*cKO^FEMALE^) (ANOVA, *p=0.019; Fig. 2H). Altogether, these results demonstrate that the loss of *Atrx* in forebrain excitatory neurons postnatally does not result in social deficits in male and female mice.

### Repetitive behaviours are not altered in Atrx-cKO^MALE^ and Atrx-cKO^FEMALE^ mice

Previous studies using autism mouse models have demonstrated that the mutant mice often present with repetitive and stereotyped behaviours (27,34–37). We tested for the presence of these repetitive and stereotyped behaviours in both *Atrx-*cKO^MALE^ and *Atrx-*cKO^FEMALE^ mice using various tests. The marble-burying assay was used to assess repetitive burying and digging by placing mice in a cage with 12 marbles and recording how many marbles were buried following a 30-minute period. Percentage of marbles buried during the marble-burying assay was not significantly different when comparing *Atrx-*cKO^MALE^ and *Atrx-*cKO^FEMALE^ mice to control mice. There also was no difference when comparing between genotypes or sexes. However, the interaction was significantly different between groups, suggesting the loss of *Atrx* in forebrain excitatory neurons has opposing effects on marble burying when comparing male and female mice (ANOVA, *p=0.047; Fig. **3A**).

We also investigated the presence of repetitive grooming tendencies by misting mice with water to induce grooming behaviours. The total amount of time spent grooming during the 30-minute induced self-grooming assay was not significantly different between *Atrx-*cKO mice and controls, or between sexes (Fig. 3B). When the results of this test were analyzed over 5-minute intervals, similarly, there was no difference in the amount of time spent grooming between *Atrx-* cKO^MALE^ and Ctrl^MALE^ mice (Fig. 3C) or *Atrx-*cKO^FEMALE^ and Ctrl^FEMALE^ mice (Fig. 3D). Interestingly, the open field test shows a significant increase in the number of vertical episodes (including rearing and jumping) of female mice (*Atrx-*cKO^FEMALE^ and Ctrl^FEMALE^ mice) compared to male mice (*Atrx-*cKO^MALE^ and Ctrl^MALE^) (ANOVA, ****p<0.0001; Fig. 3E). When these results were analyzed in 10-minute intervals, it is apparent that these sex-differences in vertical episodes occurred primarily within the first 60 minutes of the open-field test (multiple comparisons; *p<0.05, **p<0.01; Fig. 3F). In addition to these sex-differences, there was no genotypic difference between the total vertical episodes or the vertical episodes over time. Similarly, there were no significant difference between or *Atrx-*cKO mice and controls when analyzing vertical episodes. These behavioural analyses suggest that the loss of *Atrx* in forebrain excitatory neurons postnatally does not result in repetitive or stereotyped behaviours typically associated with ASDs. However, we did observe an increase in the number of vertical episodes of female mice compared to males, suggesting the presence of innate repetitive behaviours in female mice.

**Figure 3.**
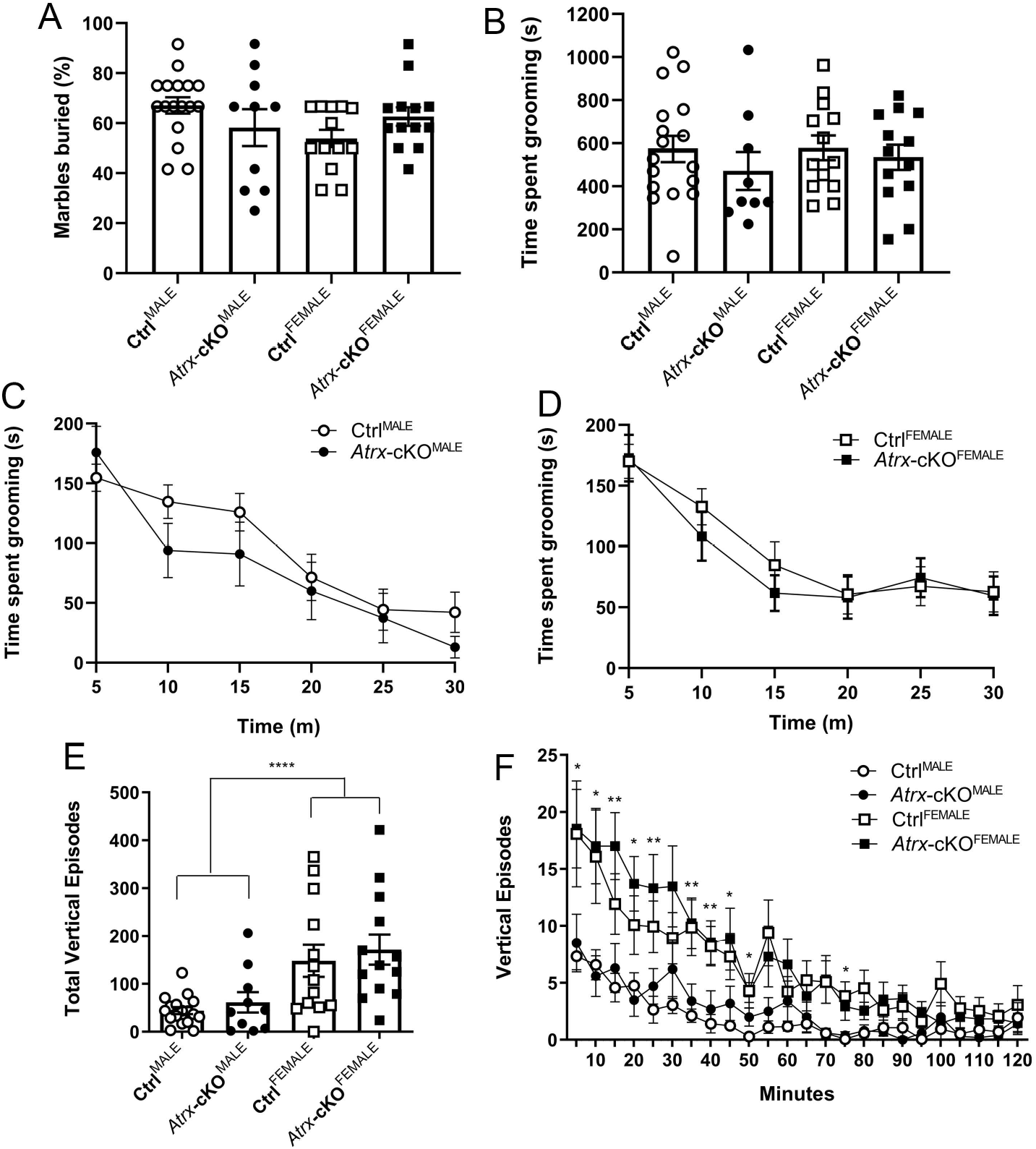
*Atrx*-cKO^MALE^ and *Atrx*-cKO^FEMALE^ mice display sex-differences, but not genotypic-differences in stereotyped behaviours. A) Percentage of marbles buried during 30-minute marble burying task (twANOVA, genotype, F_(1,49)_=0.0003, p=0.985; tw ANOVA, sex, F_(1,49)_=1.038, p=0.313; twANOVA, interaction, F_(1,49)_=4.132, *p=0.047). B) Total time grooming during 30-minute water-induced grooming task (twANOVA, genotype, F_(1,48)_=1.200, p=0.279; twANOVA, sex, F_(1,48)_=0.268, p=0.607). C-D) Amount of time spent grooming over 5-mintue intervals during water-induced grooming task (twANOVA, males, F_(1,44)_=2.386, p=0.125; twANOVA, females, F_(1,44)_=0.562, p=0.455). E) Total number of vertical episodes during 120-minute open field test (twANOVA, genotype, F_(1,49)_=0.673, p=0.416; twANOVA, sex, F_(1,49)_=18.71, ****p<0.0001). F) Number of vertical episodes over 10-minute intervals during the 120-minute open field test (tw ANOVA, genotype, F_(1,51)_=0.866, p=0.356; twANOVA, sex, F_(1,51)_=19.19, ****p<0.0001; mc, *p<0.05, **p<0.01). *Atrx*-cKO^MALE^: n=10, Ctrl^MALE^: n=17, *Atrx*-cKO^FEMALE^: n=13, Ctrl^FEMALE^: n=13. Error bars: ± SEM.

### Atrx-cKO^MALE^ and Atrx-cKO^FEMALE^ mice display typical startle response to acoustic stimuli

Previous studies have reported that rodent models with autism-associated genetic mutations can display an exaggerated startle response, or impaired pre-pulse inhibition, to an acoustic stimulus (38–41). These impairments are associated with deficits in sensory gating and auditory processing often reported in ASD patients (42,43). As such, we wanted to investigate if *Atrx*-cKO mice display hypersensitivity to an acoustic startle stimulus. The pre-pulse inhibition and startle response assay using an acoustic stimulus was performed, as described previously (31). Mice were placed in a chamber and exposed to fifty trials of an acoustic stimulus (20 ms, 115 db). *Atrx-*cKO^MALE^ and *Atrx-*cKO^FEMALE^ mice demonstrated a similar startle response to the acoustic stimuli compared to their respective controls (Fig. 4A-B). Additionally, there was no significant difference in startle responses when comparing genotypes (*Atrx-*cKO vs. Ctrl). Notably, there was a significant increase in the startle response of male mice (*Atrx-*cKO^MALE^ and Ctrl^MALE^) compared to female mice (*Atrx-*cKO^FEMALE^ and Ctrl^FEMALE^) (ANOVA, *p<0.023). We also performed a set of trials that investigated pre-pulse inhibition to the acoustic stimulus by first exposing mice to a pre-pulse that preceded the acoustic stimulus. These trials varied in the intensity of the pre-pulse (75 db or 80db), and the amount of time between the pre-pulse and the acoustic “pulse” stimulus (30ms or 100ms). Additionally, there was one trial that only involved a “pulse” without a pre-pulse. Startle responses to this “pulse-only” trial were used as a baseline, and results from all other trials were expressed as a percentage of this baseline. For all pre-pulse trials, both *Atrx-*cKO^MALE^ and *Atrx-*cKO^FEMALE^ mice did not demonstrate significant difference in their startle responses to the acoustic stimulus compared to the controls. Overall, these results suggest that the loss of *Atrx* in neurons does not result in an exaggerated startle response or impaired pre-pulse inhibition in male or in female mice. However, when analyzing these results grouped by sex, male mice display an increased startle response to an acoustic stimulus compared to female mice.

**Figure 4.**
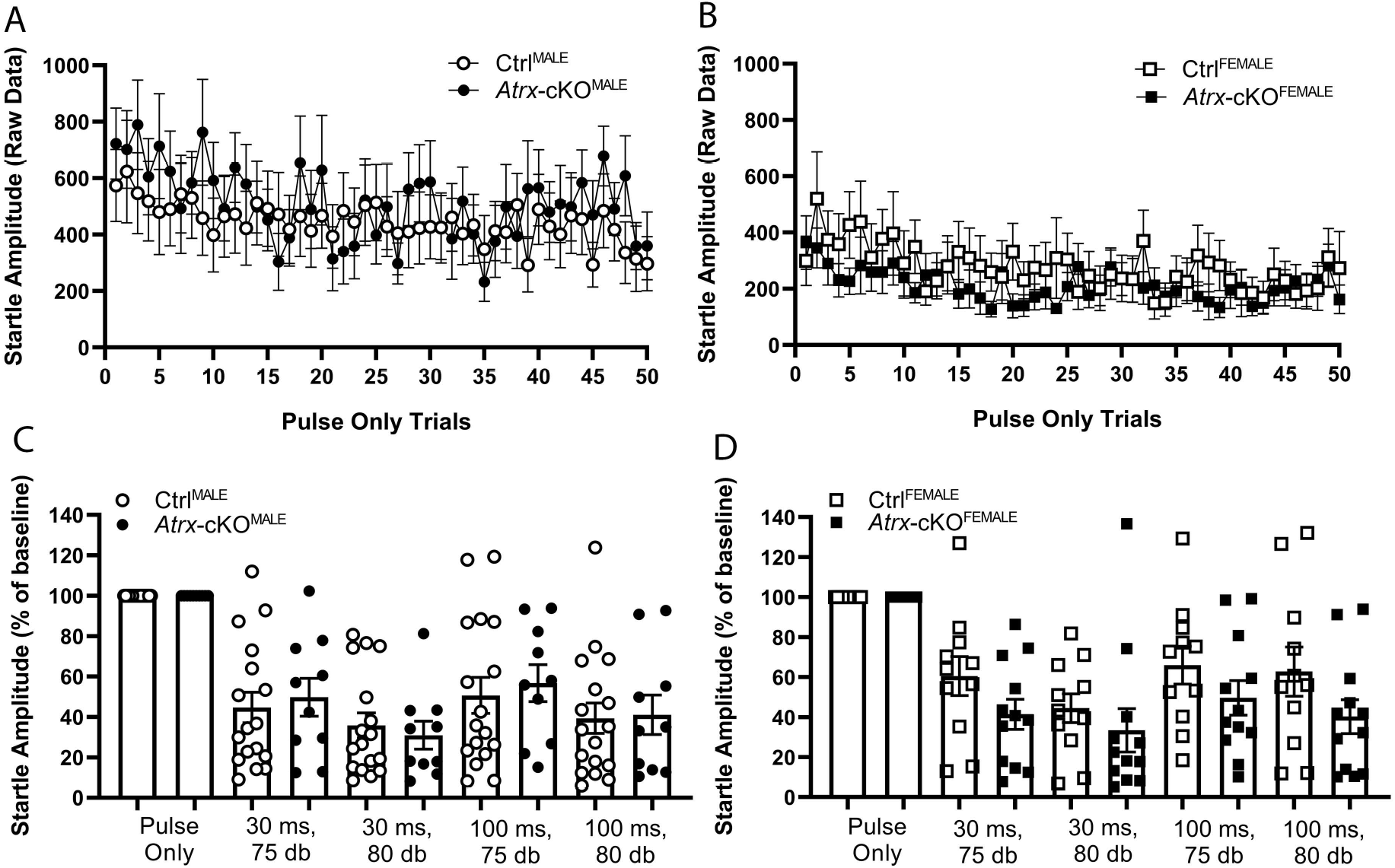
Acoustic startle response is exaggerated in *Atrx*-cKO^MALE^ and Ctrl^MALE^ mice compared to females. A-B) Quantification of startle responses (recorded in millivolts) to fifty “pulse only” trials (twANOVA, genotype, F_(1,48)_=0.084, p=0.773; twANOVA, sex, F_(1,48)_=5.555, *p=0.023). C-D) Averaged pre-pulse inhibition for four trial types, varying in interstimulus intervals (30 ms or 100 ms) and pre-pulse intensity (75 db or 80 db). Startle responses for these trials is expressed as a percentage of the normalized “pulse only” trial (baseline) (Welch’s t-test, *Atrx*-cKO^MALE^ and Ctrl^MALE^, p>0.05; Welch’s t-test, *Atrx*-cKO^FEMALE^ and Ctrl^FEMALE^, p>0.05). *Atrx*-cKO^MALE^: n=10, Ctrl^MALE^: n=17, *Atrx*-cKO^FEMALE^: n=11, Ctrl^FEMALE^: n=12. Error bars: ± SEM.

## Discussion

In this study we present a comprehensive assessment of the effects of a postnatal conditional knockout of the autism susceptibility gene *Atrx* on autistic traits in male and female mice. We provide evidence that the postnatal loss of *Atrx* in forebrain excitatory neurons does not result in social deficits, stereotypies and repetitive behaviours, or sensory gating deficits. We did, however, identify sex-specific differences in sociability, vertical episode stereotypies, and startle responses when analyses were grouped by sex.

Prior to investigating ASD-related behaviours, we first sought to determine whether *Atrx*-cKO adult mice present with any olfactory defects compared to controls. We performed the odor discrimination and habituation assay to assess if *Atrx*-cKO mice showed habituation to a repeatedly presented odor and were able to discriminate between a novel odor (25). By establishing that experimental mice are able to detect and discriminate between odors, results of subsequent social behaviour assays can be more accurately interpreted. Results of the odor habituation and discrimination assay suggest that *Atrx*-cKO^MALE^ mice have deficits in olfaction, as they spent overall less time smelling multiple odors compared to controls. In particular, odor discrimination may be affected in *Atrx*-cKO^MALE^ mice as demonstrated by decreased time smelling a novel odor following repeated presentation of another. These differences in olfaction that *Atrx*-cKO^MALE^ mice display is a variable to consider when interpreting social behaviour assays that require odor discrimination. However, it is interesting to note that olfaction deficits have been reported in both ASD patients as well as models with autism-associated genetic mutations. In a recent clinical study, nine children with ASD demonstrated impaired olfactory adaption compared to a control group (44). Additionally, another study reported that mice with haploinsufficiency of the autism-associated gene, T-box, Brain 1 (*Tbr1*), displayed impairments of olfactory discrimination (45). Therefore, these impairments in olfaction, particularly odor discrimination, may be an indication of autistic-like features.

While there was no overall difference in olfaction when we compared *Atrx-*cKO^FEMALE^ mice to controls, *Atrx-*cKO^FEMALE^ mice demonstrated increased time spent sniffing the cotton swab when presented with the banana odor for the first trial. These results suggest that their odor discrimination may be heightened compared to that of Ctrl^FEMALE^. Altogether, although *Atrx*-cKO^MALE^ and *Atrx-*cKO^FEMALE^ mice display differences in olfaction compared to controls, we did not identify any genotypic-differences in social behaviours either in the social approach test or the 3-chamber paradigm. Therefore, any differences in olfaction of *Atrx-*cKO mice did not result in impairments in social recognition or discrimination based on olfactory cues.

Although there were no identified genotypic effects on social behaviour, we did discover sex-differences in both sociability as well as social novelty. Male mice spent more time socially interacting with a stranger mouse compared to females during the social approach assay. Additionally, male mice showed a greater preference for a novel mouse over a familiar mouse during the 3-chamber paradigm, while female mice did not. This increased sociability in males has been previously observed in *C57Bl/6N* mice during social behaviour assays (46). In a study by Karlsson e*t al.,* both male and female mice showed sociability and social memory, however females showed lower interaction times towards novel conspecifics compared to males. This increased sociability was attributed to higher level testosterone in males compared to females (46). Previous research has suggested that estrogen functions to facilitate social recognition in female mice (47,48). Therefore, since our study did not include information about the estrus cycle phase of females that performed the social behaviour assays, estrogen levels may have been varying amongst females during these assays. These changes in estrogen levels may provide an explanation as to why both *Atrx-*cKO^FEMALE^ and Ctrl^FEMALE^ mice did not demonstrate a preference to the novel mouse during the social memory assay.

We additionally observed sex differences in the total number of vertical episodes, and vertical episodes over time, during the open field assay. *Atrx-*cKO^FEMALE^ and Ctrl^FEMALE^ mice exhibit more vertical episodes compared to males, particularly within the first 60 minutes of the test. Results from the open field test can be interpreted in various ways, and therefore, are more powerful when analyzed in combination. Previous work in our lab had identified additional sex-differences in the open field test, with *Atrx-*cKO^FEMALE^ and Ctrl^FEMALE^ mice spending less time in centre compared to males(21). Decreased time in centre as well as increased vertical episodes in the open field test have been previously interpreted as a demonstration of anxiety-like behaviour (49,50). Therefore, these results suggest that *Atrx-*cKO^FEMALE^ and Ctrl^FEMALE^ mice display heightened anxiety-like behaviour compared *Atrx*-cKO^MALE^ and Ctrl^MALE^ mice.

Finally, we also observed a sex-specific difference in startle responses when we exposed male and female mice to an acoustic startle. Male mice demonstrated an exaggerate startle when exposed to 50 trials of a “pulse-only” acoustic stimulus compared to females. Increased startle response of males compared to female has previously been reported in studies investigating startle of mice as well as humans (51–54). Plappert *et al*., investigated startle responses of both *C57* and *C3H* mouse strains, and observed a higher startle amplitude for males, irrespective of strain. This study suggested that the estrous cycle of females did not influence acoustic startle amplitude. Rather, increased anxiety in male mice appeared to contribute to this increase in startle response (53). Interestingly, results from our open field test suggest that *Atrx-*cKO^FEMALE^ and Ctrl^FEMALE^ mice exhibit increased anxiety-like behaviours. Therefore, our results instead suggest that increased anxiety of female mice may limit their response to an acoustic startle. An alternative explanation of these sex-differences may be that *Atrx*-cKO^MALE^ and Ctrl^MALE^ mice weigh more than their female counterparts, resulting in a greater startle amplitude measured by the recording platform when the acoustic startle is administered.

Altogether, although specific sex-difference were observed throughout the tests, there were no genotypic differences between control mice and *Atrx*-cKO mice. We theorize that the lack of autistic-like phenotypes observed in *Atrx*-cKO mice is due to the timing at which *Atrx* is deleted in forebrain excitatory neurons (starting at postnatal day 18-20). ASD is clinically defined as a developmental disorder due to the majority of symptoms becoming apparent in the first few years of life. As such, genetic mutations that contribute to autistic phenotypes may need to occur during embryogenesis or be inherited (1,2,4). Future studies should utilize additional Cre/loxp systems to investigate if the loss of *Atrx* in differentiated forebrain excitatory neurons during embryogenesis results in autistic-like behaviours in male and female mice.

## Conclusions

In conclusion, a postnatal conditional knockout of the autism susceptibility gene *Atrx* did not result in autistic traits in either male or female mice. We did observe sex-specific differences in sociability, vertical episodes, and acoustic startle response that support previously reported findings. These findings suggest that the postnatal loss of ASD associated genes may occur too late in development to result in behavioural deficits. Our results support the belief that ASD is a developmental disorder and associated genetic mutations must occur early in development to result in behavioural differences.

## Abbreviations

Atrx: Alpha-thalassemia mental retardation, X-linked
cKO: conditional knockout
ASD: autism spectrum disorder
twANOVA: two-way ANOVA
mc: multiple comparisons

## Declarations

### Ethics approval and consent to participate

Not applicable

### Consent for publication

Not applicable

### Availability of data and materials

All data generated or analysed during this study are included in this published article.

### Authors' contributions

NMK participated in the design of the study, carried out the experiments and genotyping, analyzed the data, interpreted the results, and wrote the manuscript. NGB contributed to the conception and design of the study, interpretation of data, and writing of the article. All authors read and approved the final manuscript.

## Acknowledgements

We are grateful to Doug Higgs and Richard Gibbons for the Atrx floxed mice, and Vania Prado and Marco Prado for the CaMKII-Cre mice. We are also grateful to Renee Tamming and Radu Gugustea for their aid in mouse husbandry. All behavioural tests were performed at the Robarts Research Institute neurobehavioural core facility.

## References

1. Abrahams BS, Geschwind DH. Advances in autism genetics: On the threshold of a new neurobiology. Nat Rev Genet. 2008;9(5):341–55.

2. Chen JA, Peñagarikano O, Belgard TG, Swarup V, Geschwind DH. The Emerging Picture of Autism Spectrum Disorder: Genetics and Pathology. Annu Rev Pathol Mech Dis. 2015;10(1):111–44.

3. Sanders SJ, Murtha MT, Gupta AR, Murdoch JD, Raubeson MJ, Willsey J, et al. De novo mutations revealed by whole exome sequencing are strongly associated with autism. Nature. 2013;485(7397):237–41.

4. Guo H, Hu Z, Zhao J, Xia K. Genetics of autism spectrum disorders. J Cent South Univ (Medical Sci. 2011;36(8):703–11.

5. Yang G, Sau C, Lai W, Cichon J, Li W. Synaptic, transcriptional, and chromatin genes disrupted in autism. 2015;344(6188):1173–8.

6. Loke YJ, Hannan AJ, Craig JM. The role of epigenetic change in autism spectrum disorders. Front Neurol. 2015;6:1–18.

7. Silverman JL, Yang M, Lord C, Crawley JN. Behavioural phenotyping assays for mouse models of autism. Nat Rev Neurosci. 2010;11(7):490–502.

8. Crawley JN. Behavioral Phenotyping Strategies for Mutant Mice. Neuron. 2008;57(6):809–18.

9. Gibbons RJ, Picketts DJ, Villard L, Higgs DR. Mutations in a putative global transcriptional regulator cause X-linked mental retardation with α-thalassemia (ATR-X syndrome). Cell. 1995;80(6):837–45.

10. Gibbons RJ, Wada T, Fisher CA, Malik N, Mitson MJ, Steensma DP, et al. Mutations in the chromatin-associated protein ATRX. Hum Mutat. 2008;29(6):796–802.

11. Gibbons R. Alpha thalassaemia-mental retardation, X linked. Orphanet J Rare Dis. 2006;1(1):1–9.

12. Sun M, Paciga JE, Feldman RI, Yuan ZQ, Coppola D, You Yong Lu, et al. Large-scale discovery of novel genetic causes of developmental disorders. Cancer Res. 2001;61(16):5985–91.

13. Yu TW, Chahrour MH, Coulter ME, Jiralerspong S, Ataman B, Harmin D a, et al. Using whole exome sequencing to identify inherited causes of autism. 2013;77(2):259–73.

14. Gong X, Bacchelli E, Blasi F, Toma C, Betancur C, Chaste P, et al. Analysis of X chromosome inactivation in autism spectrum disorders. Am J Med Genet Part B Neuropsychiatr Genet. 2008;147(6):830–5.

15. Brett M, McPherson J, Zang ZJ, Lai A, Tan ES, Ng I, et al. Massively parallel sequencing of patients with intellectual disability, congenital anomalies and/or autism spectrum disorders with a targeted gene panel. PLoS One. 2014;9(4):1–9.

16. Munnich A, Demily C, Frugère L, Duwime C, Malan V, Barcia G, et al. Impact of on-site clinical genetics consultations on diagnostic rate in children and young adults with autism spectrum disorder. Mol Autism. 2019;10(1):1–10.

17. Aspromonte MC, Bellini M, Gasparini A, Carraro M, Bettella E, Polli R, et al. Characterization of intellectual disability and autism comorbidity through gene panel sequencing. Hum Mutat. 2019;40(9):1346–63.

18. Li J, Wang L, Guo H, Shi L, Zhang K, Tang M, et al. Targeted sequencing and functional analysis reveal brain-size-related genes and their networks in autism spectrum disorders. Mol Psychiatry. 2017;22(9):1282–90.

19. Levy MA, Kernohan KD, Jiang Y, Bérubé NG. ATRX promotes gene expression by facilitating transcriptional elongation through guanine-rich coding regions. Hum Mol Genet. 2014;24(7):1824–35.

20. Kernohan KD, Jiang Y, Tremblay DC, Bonvissuto AC, Eubanks JH, Mann MRW, et al. ATRX Partners with Cohesin and MeCP2 and Contributes to Developmental Silencing of Imprinted Genes in the Brain. Dev Cell. 2010;18(2):191–202.

21. Tamming R, Dumeaux V, Langlois L, Ellegood J, Qiu LR, Jiang Y, et al. Atrx deletion in neurons leads to sexually-dimorphic dysregulation of miR-137 and spatial learning and memory deficits. bioRxiv. 2019;606442.

22. Watson LA, Solomon LA, Li JR, Jiang Y, Edwards M, Shin-Ya K, et al. Atrx deficiency induces telomere dysfunction, endocrine defects, and reduced life span. J Clin Invest. 2013;123(5):2049–63.

23. Bérubé NG, Mangelsdorf M, Jagla M, Vanderluit J, Garrick D, Gibbons RJ, et al. The chromatin-remodeling protein ATRX is critical for neuronal survival during corticogenesis. J Clin Invest. 2005;115(2):258–67.

24. Tsien JZ, Chen DF, Gerber D, Tom C, Mercer EH, Anderson DJ, et al. Subregion- and cell type-restricted gene knockout in mouse brain. Cell. 1996;87(7):1317–26.

25. Arbuckle EP, Smith GD, Gomez MC, Lugo JN. Testing for Odor Discrimination and Habituation in Mice. J Vis Exp. 2015;(99):1–7.

26. Jamain S, Radyushkin K, Hammerschmidt K, Granon S, Boretius S, Varoqueaux F, et al. Reduced social interaction and ultrasonic communication in a mouse model of monogenic heritable autism. Proc Natl Acad Sci. 2008;105(5):1710–5.

27. El-Kordi A, Winkler D, Hammerschmidt K, Kästner A, Krueger D, Ronnenberg A, et al. Development of an autism severity score for mice using Nlgn4 null mutants as a construct-valid model of heritable monogenic autism. Behav Brain Res. 2013;251:41–9.

28. Deacon RMJ. Digging and marble burying in mice: simple methods for in vivo identification of biological impacts. 2006.

29. Kalueff A V., Wayne Aldridge J, Laporte JL, Murphy DL, Tuohimaa P. Analyzing grooming microstructure in neurobehavioral experiments. Nat Protoc. 2007;2(10):2538–44.

30. Tamming RJ, Siu JR, Jiang Y, Prado MAM, Beier F, Bérubé NG. Mosaic expression of *Atrx* in the mouse central nervous system causes memory deficits. Dis Model Mech. 2017;10(2):119–26.

31. Valsamis B, Schmid S. Habituation and Prepulse Inhibition of Acoustic Startle in Rodents. J Vis Exp. 2011;(55):1–10.

32. Tong D li, Chen R guo, Lu Y lan, Li W ke, Zhang Y fang, Lin J kai, et al. The critical role of ASD-related gene CNTNAP3 in regulating synaptic development and social behavior in mice. Neurobiol Dis. 2019;130:104486.

33. Courchet V, Roberts AJ, Meyer-Dilhet G, Del Carmine P, Lewis TL, Polleux F, et al. Haploinsufficiency of autism spectrum disorder candidate gene NUAK1 impairs cortical development and behavior in mice. Nat Commun. 2018;9(1).

34. Nuytens K, Gantois I, Stijnen P, Iscru E, Laeremans A, Serneels L, et al. Haploinsufficiency of the autism candidate gene Neurobeachin induces autism-like behaviors and affects cellular and molecular processes of synaptic plasticity in mice. Neurobiol Dis. 2013;51:144–51.

35. Burrows EL, Laskaris L, Koyama L, Churilov L, Bornstein JC, Hill-Yardin EL, et al. A neuroligin-3 mutation implicated in autism causes abnormal aggression and increases repetitive behavior in mice. Mol Autism. 2015 Nov 14;6(1).

36. Cheng Y, Wang ZM, Tan W, Wang X, Li Y, Bai B, et al. Partial loss of psychiatric risk gene Mir137 in mice causes repetitive behavior and impairs sociability and learning via increased Pde10a. Nat Neurosci. 2018;21.

37. Jung H, Park H, Choi Y, Kang H, Lee E, Kweon H, et al. Sexually dimorphic behavior, neuronal activity, and gene expression in Chd8-mutant mice. Nat Neurosci. 2018;21(9):1218–28.

38. Jiang DY, Wu Z, Forsyth CT, Hu Y, Yee SP, Chen G. GABAergic deficits and schizophrenia-like behaviors in a mouse model carrying patient-derived neuroligin-2 R215H mutation. Mol Brain. 2018;11(1):1–11.

39. Luo J, Norris RH, Gordon SL, Nithianantharajah J. Neurodevelopmental synaptopathies: Insights from behaviour in rodent models of synapse gene mutations. Vol. 84, Progress in Neuro-Psychopharmacology and Biological Psychiatry. Elsevier Inc.; 2018. p. 424–39.

40. Crawley JN. Mouse behavioral assays relevant to the symptoms of autism. Brain Pathol. 2007;17(4):448–59.

41. Scott KE, Schormans AL, Pacoli K, De Oliveira C, Allman BL, Schmid S. Altered auditory processing, filtering, and reactivity in the *Cntnap2* knockout rat model for neurodevelopmental disorders. J Neurosci. 2018;0759–18.

42. Takahashi H, Kamio Y. Acoustic startle response and its modulation in schizophrenia and autism spectrum disorder in Asian subjects. Schizophr Res. 2018;198:16–20.

43. Ebishima K, Takahashi H, Stickley A, Nakahachi T, Sumiyoshi T, Kamio Y. Relationship of the acoustic startle response and its modulation to adaptive and maladaptive behaviors in typically developing children and those with autism spectrum disorders: A pilot study. Front Hum Neurosci. 2019;13:1–6.

44. Kumazaki H, Okamoto M, Yoshimura Y, Ikeda T, Hasegawa C, Saito DN, et al. Brief Report: Odour Awareness in Young Children with Autism Spectrum Disorders. J Autism Dev Disord. 2018.

45. Huang TN, Yen TL, Qiu LR, Chuang HC, Lerch JP, Hsueh YP. Haploinsufficiency of autism causative gene Tbr1 impairs olfactory discrimination and neuronal activation of the olfactory system in mice. Mol Autism. 2019;10(1):1–16.

46. Karlsson SA, Haziri K, Hansson E, Kettunen P, Westberg L. Effects of sex and gonadectomy on social investigation and social recognition in mice. BMC Neurosci. 2015;16(1):1–10.

47. Tang AC, Nakazawa M, Romeo RD, Reeb BC, Sisti H, McEwen BS. Effects of long-term estrogen replacement on social investigation and social memory in ovariectomized C57BL/6 mice. Horm Behav. 2005;47(3):350–7.

48. Sánchez-Andrade G, Kendrick KM. Roles of α- and β-estrogen receptors in mouse social recognition memory: Effects of gender and the estrous cycle. Horm Behav. 2011;59(1):114–22.

49. Borta A, Schwarting RKW. Inhibitory avoidance, pain reactivity, and plus-maze behavior in Wistar rats with high versus low rearing activity. Physiol Behav. 2005;84(3):387–96.

50. Seibenhener ML, Wooten MC. Use of the open field maze to measure locomotor and anxiety-like behavior in mice. J Vis Exp. 2015;(96):1–6.

51. Kofler M, Müller J, Reggiani L, Valls-Solé J. Influence of gender on auditory startle responses. Brain Res. 2001;921(1-2):206–10.

52. Aasen I, Kolli L, Kumari V. Sex effects in prepulse inhibition and facilitation of the acoustic startle response: Implications for pharmacological and treatment studies. J Psychopharmacol. 2005;19(1):39–45.

53. Plappert CF, Rodenbücher AM, Pilz PKD. Effects of sex and estrous cycle on modulation of the acoustic startle response in mice. Physiol Behav. 2005;84(4):585–94.

54. Kumari V, Aasen I, Papadopoulos A, Bojang F, Poon L, Halari R, et al. A comparison of prepulse inhibition in pre- and postmenopausal women and age-matched men. Neuropsychopharmacology. 2008;33(11):2610–8.

